# Simulation of P2X-mediated calcium signaling in microglia

**DOI:** 10.1101/354142

**Authors:** Ben Chun, Bradley D. Stewart, Darin Vaughan, Adam D. Bachstetter, Peter M. Kekenes-Huskey

## Abstract

Microglia function is orchestrated through highly-coupled signaling pathways that depend on calcium (Ca^2+^). In response to extracellular adenosine triphosphate (ATP), transient increases in intracellular Ca^2+^ driven through the activation of purinergic receptors, P_2_X and P_2_Y, are sufficient to promote cytokine synthesis and potentially their release. While steps comprising the pathways bridging purinergic receptor activation with transcriptional responses have been probed in great detail, a quantitative model for how these steps collectively control cytokine production has not been established. Here we developed a minimal computational model that quantitatively links extracellular stimulation of two prominent ionotropic purinergic receptors, P_2_X_4_ and P_2_X_7_, with the graded production of a gene product, namely the tumor necrosis factor *α* (TNF*α*) cytokine. In addition to Ca^2+^ handling mechanisms common to eukaryotic cells, our model includes microglia-specific processes including ATP-dependent P_2_X_4_ and P_2_X_7_ activation, activation of NFAT transcription factors, and TNF*α* production. Parameters for this model were optimized to reproduce published data for these processes, where available. With this model, we determined the propensity for TNF*α* production in microglia, subject to a wide range of ATP exposure amplitudes, frequencies and durations that the cells could encounter *in vivo.* Furthermore, we have investigated the extent to which modulation of the signal transduction pathways influence TNF*α* production. Our key findings are that TNF*α* production via P_2_X_4_ is maximized at low ATP when subject to high frequency ATP stimulation, whereas P_2_X_7_ contributes most significantly at millimolar ATPranges. Given that Ca^2+^ homeostasis in microglia is profoundly important to its function, this computational model provides a quantitative framework to explore hypotheses pertaining to microglial physiology.

## 1 Introduction

Microglia, the tissue macrophages of the central nervous system (CNS), account for approximately 10% of CNS cells. Beyond their role as immune cells, microglia have essential physiological functions that maintain the health of the nervous system. Microglia express cell surface receptors, which, in ways that are not fully understood, integrate extracellular signals in the milieu to guide microglia to modify their local environment selectively. Examples of how microglia modulate neuronal circuits to alter CNS function include the pruning of synapses (for review see: [1]), and through the release of bioactive molecules, such as cytokines. The cytokine tumor necrosis fa+ctor alpha (TNF*α*) is a microglia secreted molecule that can directly increase and decrease the balance of excitatory and inhibitory neurotransmission, by regulating AMPA and GABAa receptors (for review see: [2]). In the bidirectional communication between microglia, and other neural cells, ATP has a particularly significant role in the cell-to-cell communication of microglia. While some of the microglia signal transduction pathways activated by ATP are known in considerable detail (for review see: [3]), how these pathways are integrated and scaled to orchestrate a specific cellular response remains to be defined. Computational systems biology has emerged as a powerful tool towards bridging external stimuli (i.e.,ATP) with phenotypical outputs (i.e., TNF*α*)[4]. Our goal is to establish a computational systems biology model of ATP-dependent signaling pathways in microglia to probe for factors that contribute to increased TNF*α*. To our knowledge, models for ATP-dependent glial function are scarce. Some limited modeling of purinergic receptor gating (P_2_X)[5, 6, 3, 7, 8, 9, 10, 11], as well as IP_3_-mediated Ca^2+^ signaling[12] has been generated, but these models have not integrated the signaling mechanisms starting with external ATP stimuli and ultimately leading to TNF*α*. ATP binds to purinoceptors localized to microglial plasma membrane (PM). Purinoceptor activation is strongly associated with dramatic changes in intracellular Ca^2+^ content and Ca^2+^-dependent signaling cascades linked to gene expression. Changes in Ca^2+^ in response to ATP can occur directly through the opening of Ca^2+^ non-selective ionotropic channels (P_2_X), or indirectly by metabotropic receptors (P_2_Y) that utilize second messengers like IP_3_. Of the P_2_X receptors, P_2_X_4_ and P_2_X_7_ are the most abundant channels expressed in microglial membrane[3]. These channels respond to micromolar and millimolar ATP, respectively,[7, 13] with P_2_X_4_ generally presenting more rapid deactivation times compared to P_2_X_7_[14]. P_2_X activation by ATP causes increased production and release of TNF*α* by microglia [8]. ATP-dependent TNF*α* production is also strongly correlated with both Ca^2+^ and increased p38 phosphoylation[8]. Ca^2+^-dependence of NFAT activation via calcineurin[15,16] has also been implicated in the ATP-dependent release of TNF*α* from microglia[17,18].

We hypothesized that the increased microglia TNF*α* production, controlled dynamically by ATP binding to P_2_X receptors, involves Ca^2+^ signaling and namely calcineurin (CN)-nuclear factor of activated T-cells (NFAT) signal transduction. Toward investigating this hypothesis, we integrated Markov models of P_2_X_4_ and P_2_X_7_ activation[5, 19], and Ca^2+^ homeostasis[15, 20, 21, 22].
Microglia utilize diverse mechanisms to control Ca^2+^ homeostasis and Ca^2+^-dependent responses to external stimuli, of which, many are broadly used by nearly all eukaryotic cells. These mechanisms encompass external Ca^2+^ entry, the triggering of intracellular Ca^2+^ release by way of second messengers, Ca^2+^ binding to buffers and effectors, as well as Ca^2+^ clearance via pumps and exchangers[23]. The mechanisms of Ca^2+^ homeostasis tested in our model include Na+/Ca^2+^exchanger (NCX) and Sarcoplasmic/endoplasmic reticulum calcium ATPase (SERCA), and Ca^2+^-dependent control of TNF*α* transcription, for which models have been developed for other cell types, such as cardiac cells [21, 24]. Results of our computational model suggest that short/pulsatile ATP is likely sufficient to invoke production of the TNF*α* via P_2_X_4_. Long duration and high amplitude ATP exposure is necessary for production via P_2_X_7_. Both effects are determined critically by sustained elevation of Ca^2+^ load that exceed intrinsic rates of NCX-mediated Ca^2+^ extrusion. In summation, with this computational model, we provide a framework to challenge our current understanding of microglia physiology and motivate new hypotheses and avenues of experiments.

## 2 Methods

**Figure 1:**
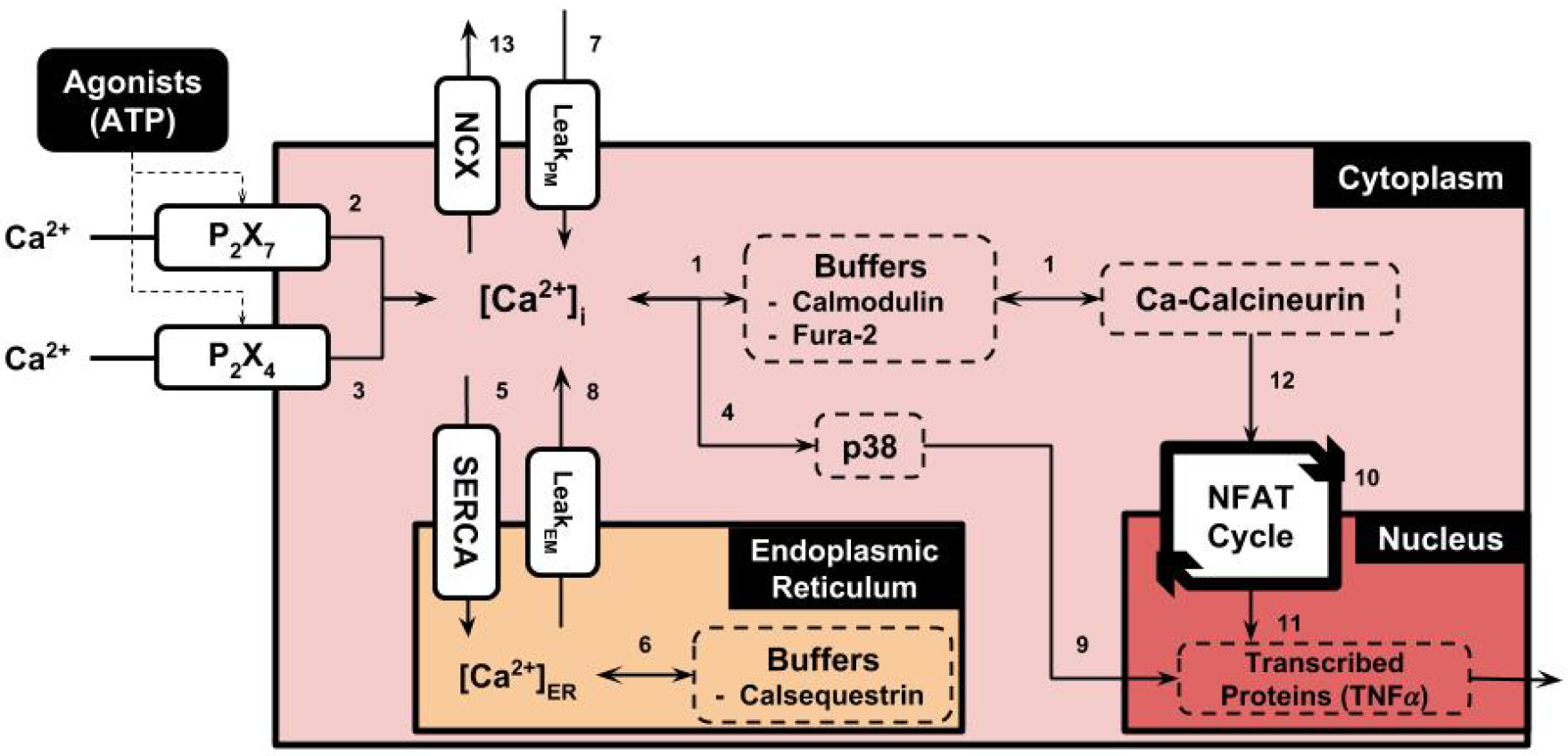
Schematic of the microglial signaling model assumed in this study. Table S1 provides a summary of the numbered pathways in this diagram, as well as a link to the corresponding equations in the Supplement. In this schematic, ATP activates the purinoceptors, P_2_X_4_and P_2_X_7_, which culminates in the exocytosis of TNF*α*.

### 2.1 Microglial Ca^2+^ model (Fig. 1)

Our computational model of P_2_X-mediated microglial signaling portrayed in Fig. 1 assumes:

1. ATP-dependent activation of P_2_X receptors, assuming pharmacologic inhibition of P_2_Y receptors [25] (see Sect. A.3.1)
2. Ca^2+^-dependent activation of signaling proteins calmodulin (CaM), CN, NFAT, as well as Ca^2+^ extrusion and uptake mechanisms (see Sect. A.3.4)
3. TNF*α* transcription, translation and exocytosis (see Sect. A.3.5). For clarity, we numbered these reactions and list their supporting literature in Table S1. While microglial regulatory mechanisms are complex and involve a great variety of channels, proteins and processes that we neglect here, our minimal model permits quantitative descriptions of P_2_X-mediated behavior with a tractable number of equations and parameters. These equations form the basis of time-dependent ordinary differential equations (see Sect. A.3) that we numerically integrate, subject to a variety of experimental conditions described below. Parameters for these equations are summarized in Sect. A.1. Where possible, these parameters were adopted from the literature, although in many cases these parameters were refitted to reproduce experimental data measured in microglia. The experimental data against which our models were fitted or validated were drawn from pubmed database searches. If appropriate, we identified in the validation commentary data sets that were in disagreement. Frequently, variation in data sets could be attributed to differences in cell lines used by the experimentalists, conditions to which the cells were subjected, as well as known heterogeneities in cell phenotype [26, 27]. Since cell-to-cell variation can be significant, we present sensitivity analyses in our results that indicate the extent to which differences in protein expression and activity influence measured observables.

### 2.2 Simulation

#### 2.2.1 Numerical model of Ca^2+^ handling

The microglial model was implemented in the Generalized ODE Translator go-tran (https://bitbucket.org/johanhake/gotran) framework as described previously [28]. The gotran model was compiled into a Python module to make use of our Python-based routines for simulation and analyses. In our numerical experiments, the microglia model was numerically integrated by the scipy function ODEINT, which utilizes the LSODA algorithm for stiff ordinary differential equations (ODE)s [29] The numerical model was integrated using a timestep of 0.1 ms for a total simulation time of up to 180 min. These simulations provide as output the time-dependent values of model states, such as intracellular Ca^2+^ or the open gates of the P_2_X channels. Model fitting proceeded by a genetic algorithm (reviewed in [30]) that iteratively improved parameter values, such as P_2_X_4_ conductance and PM Ca^2+^ leak over several generations of progeny. Parameters for the model components are summarized in Sect. A.1. Based on these parameterizations, our key model outputs were Ca^2+^ transients with respect to ATP exposure durations and concentrations, as well subsequent changes in NFAT activation and TNF*α* exocytosis. Experimentally-measured outputs, such as Ca^2+^ transient decay time and amplitude, were measured for each of the progeny; those that reduced output error relative to the experimentally-measured equivalent were stored for future generations. To validate our implementation, we present comparisons against predicted channel current profiles, intracellular Ca^2+^ content, activation of NFAT and pp38, as well as TNF*α* in the Results section.

#### 2.2.2 Analysis

To examine potential mechanisms that link increased P_2_X stimulation to TNF*α* production, we monitored and identified prominent changes in key ‘state’ variables (including total amount of calcineurin, calmodulin, nuclear factor of activated T-cells, the density of purinoceptors, and the influx capacity) relative to control conditions. These sensitivity analyses were performed to determine relative correlations between model input parameters and predicted outputs (see Supplement). Data processing was performed using SCIPY and the IPython notebooks. All codes written in support of this publication will be publicly available at https://bitbucket.org/pkhlab/pkh-lab-analyses. Simulation-generated data are available upon request.

## 3 Results

### 3.1 plasma membrane (PM) Ca^2+^ influxes and effluxes

In analogy to studies of P_2_X function in microglia conducted by Davalos *et al.* [25], for which P_2_Y activity was pharmacologically inhibited, our study assumed P_2_X channels comprise the primary mechanism of direct (ionotropic) Ca^2+^ entry following ATP treatment. We further assumed P_2_X-mediated Ca^2+^ entry arises predominantly from P_2_X_7_ and to a lesser extent, P_2_X_4_. The activation of the P_2_X_7_ and P_2_X_4_ receptors has been well-characterized in the literature, particularly through whole-cell electrophysiological methods using channels expressed in HEK cells as well as in microglia [31, 32, 33, 34,14, 35]. P_2_X_4_ activation is triggered at micromolar ATP, which culminates in a rapid (< 1 second) sharply peaked current profile that decays within tens of seconds [36]. P_2_X_7_ receptors, on the other hand, respond to millimolar ATP concentrations, but in contrast to P_2_X_4_, maintain significant current over long durations of ATP exposure [8, 37, 38]. As a first step toward the validation of our microglial Ca^2+^ model, we sought to reproduce the unique current profiles for the respective channels over appropriate ranges and durations of ATP exposure, using a Markov model adapted to microglia P_2_X activation [37].

The Markov model framework assumed in this study captures the time-evolution of channel gating states; of these states, the channel current is proportional to the probability of the open state. While Markov models have been reported in the literature[37,5,19, 39], including 12-and 13-state models from Khadra *et al.* and Zemkova *et al.*, for the sake of parsimony we merged gating states that were in rapid equilibrium relative to state transitions that abide much slower kinetics [40]. In Sect. A.3.1, we summarize the Markov models and their parameterizations for P_2_X_4_ and P_2_X_7_, respectively. Given the model reformulation, it was essential to ensure that our model reproduced experimentally-measured currents attributed to P_2_X_7_ and P_2_X_4_ in microglia. Here, we validate the fitted P_2_X_4_ and P_2_X_7_ models by comparing our predicted current profiles against those measured for P_2_X_4_ by Toulme *et al.*[9] and P_2_X_7_ by Chessell *et al.*[38].

We first present in Fig. 2A for P_2_X_4_ predicted currents versus recordings conducted using P_2_X_4_ transfected into microglia [9]. According to their study, the mutated P_2_X_4_ channels were expressed and induced 30-fold stronger inward currents compared to its normal resting cells. For this reason, the P_2_X_4_ responses assumed in our model are more typical of activated microglia [36]. In Fig. 2A we present two sets of modeling predictions: 1) using the ‘full’ model from Zemkova *et al.*and 2) our ‘lumped’ model, for which the open states Q1 and Q2 in the ‘full’ model are consolidated into a single Q12 ‘macro-state’. At 100 μM, both the full and lumped models closely follow the currents measured by Toulme *et al.*, namely, they capture the sharp peak of approximately -375 pA in the early phase of ATP pulse, after which the current rapidly decays. A notable distinction between the two model formulations is that the lumped model presents a modestly faster rate of current decay in the presence of ATP, relative to the full model. Moreover, upon removal of ATP, the lumped model current rapidly extinguishes, whereas the full model more gradually decays. In Fig. S2 we demonstrate that our Q12 ‘macro-state’ mirrors the summation of the Q1 and Q2 states with exception to the phase after ATP removal, hence the merging of the Q1 and Q2 states introduces some error into the current dynamics. Nevertheless, both model representations yield nearly identical channel current profiles for all but the final seconds of ATP treatment, thus our reduced model is sufficiently accurate for modeling rates of Ca^2+^ entry following ATP-dependent P_2_X_4_ activation. We also present current predictions at 10 μM and 1 mM ATP for comparison; as anticipated, the reduced ATP treatment leads to a smaller (125 pA) current maximum, while a 10-fold increase in ATP is indistinguishable from that reported for 100 μM ATP. Lastly, we demonstrate that the lumped model retains the desensitization behavior established in P_2_X_4_, as the channel peak amplitude is reduced following a second ATP treatment.

In Fig. 2B, we provide analogous current profiles stemming from P_2_X_7_ activation, subject to 1 × 10^−1^mM-1.0mM ATP for 0.5 seconds as reported by Chessell et *al.*[38]. In contrast to P_2_X_4_, the P_2_X_7_ current profile exhibits a slower rise to peak current compared to P_2_X_4_and a slower rate of current decay. We additionally find that the both the full and lumped models overestimate the rate at which peak current is exhibited, as well as underestimate the rate of current decay. While we could expect better agreement by introducing additional states, the high degree of overlap between the model predictions and the measured data suggests that minor discrepancy will not appreciable impact our estimates of Ca^2+^ entry by these channels. We also report in Fig. 2B the P_2_X_7_ channel profile when subject to 1.0 mM ATP over a 30 second interval. Under this condition, the channel exhibits a long-lived plateau that is sustained over the entire duration of ATP treatment. This extended plateau is a common gating attribute of P_2_X_7_ receptors [14] and is speculated to arise from a dilated open-pore structure following extended activation [35, 8]. Overall, these data suggest that our implementation of the P_2_X_4_ and P_2_X_7_ Markov models faithfully reproduce Ca^2+^ current profiles across a variety of ATP applications, and moreover, the channel dynamics exhibit vary considerably depending on the amplitude and duration of ATP exposure. Unless otherwise noted, all further applications of our model assume both channels are active and can be simultaneously activated by applied ATP.

**Figure 2:**
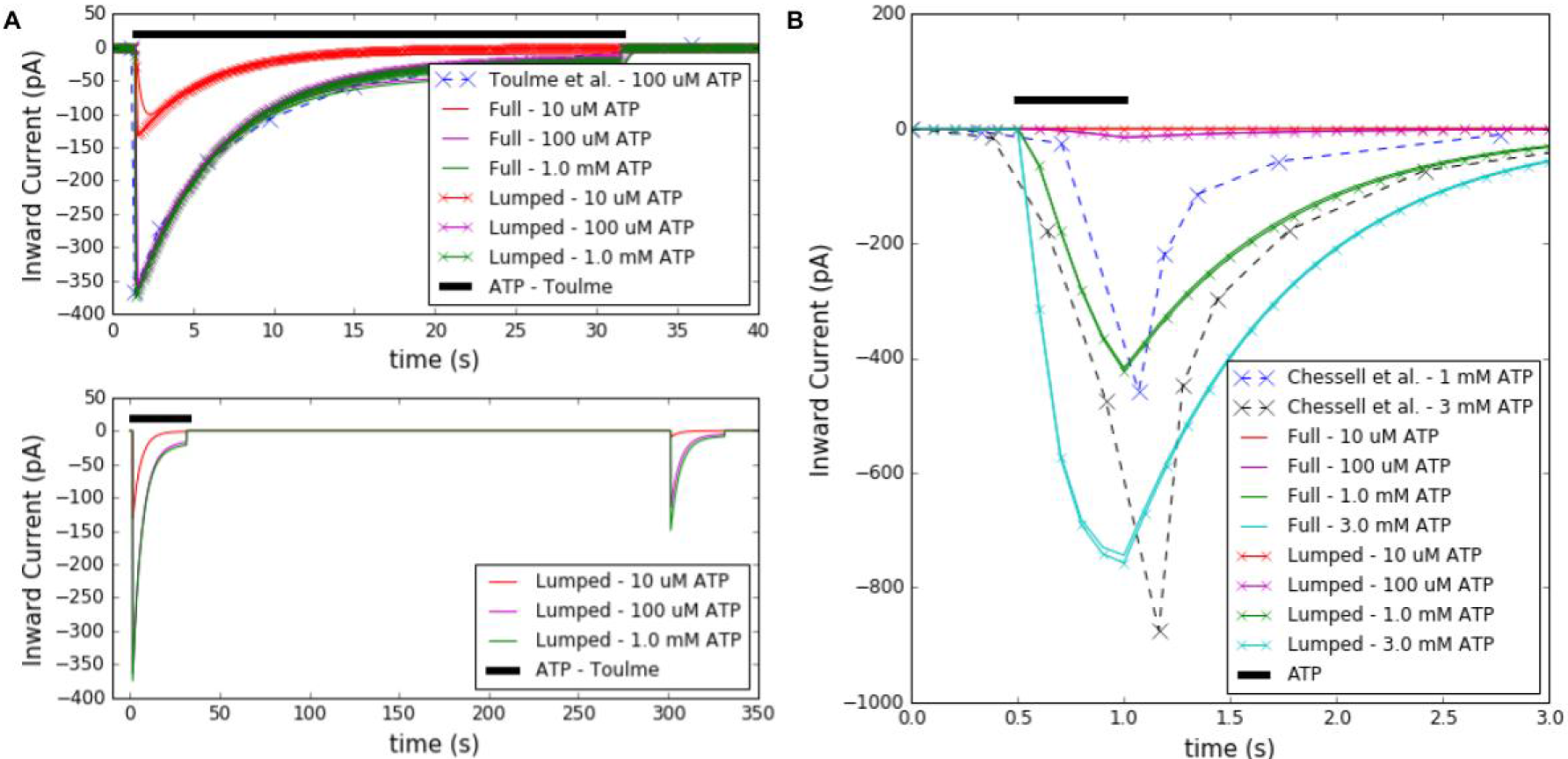
A. Inward P_2_X_4_ current profile recorded in microglia (blue dashed) by Toulme *et al.*[9] and the model predict ion (purple solid) when subject to 100 μM ATP for 30 seconds. Additional predicted profiles are given for 10 μMand 1 mM ATP to demonstrate desensitization. B. Inward P_2_X_7_ currents measured in microglia by Chessell *et al.*[38]as well as the model predictions, when subject to 10 μM to 3 mM ATP for 0.5 seconds. Parameters and conditions to model the experimental conditions are listed in Table S2.

In complement to P_2_X-driven Ca^2+^ entry, Na/Ca exchanger (NCX) and the plasma membrane ATPase comprise the primary mechanisms for extruding cytosolic Ca^2+^ and restoring basal Ca^2+^ content in glia [41, 42] and likely in microglia [3]. Given the lack of published experimental data for the role of plasma membrane ATPases in microglia Ca^2+^ management [3], we neglected their contribution in the model. In addition, a modest PM leak term was incorporated to maintain Ca^2+^ steady state at rest. This addition was necessary to counterbalance a slight NCX Ca^2+^ efflux owing to the -23 mV membrane potential assumed in the model [43]. We present in Sect. A.3.2 predictions using an NCX a model developed for NCX ion exchange in cardiac cells of rabbit origin [21], refitted here for microglia Ca^2+^ handling. Since the membrane potential in microglia under normal conditions maintains a modest negative potential of approximately -20 mV [3], NCX primarily extrudes Ca^2+^. Nevertheless, we validated our model against single-cell traces reported in Fig 5 of [43], for which significant positive NCX currents were reported from -40 mV to 60 mV (0-200 pA, see Fig. S3). Here we note that the activity characterized in [43] was conducted in Krebs medium solution, the Na+ concentration of which was comparable to the 145 mM assumed in our model. Importantly, our data successfully reproduce the experimentally measured currents under the aforementioned solution conditions. We note that our model may overestimate the contribution of NCX to Ca^2+^ extrusion rates, given the omission of PM ATPase activity. Nevertheless, our parameterization appeared sufficient to maintain resting Ca^2+^ levels, as well as restore Ca^2+^ levels following P_2_X_4_ stimulation. We illustrate this with Ca^2+^ transients reported in Fig. **??**, which demonstrate consistent relaxation to resting Ca^2+^ concentrations of 9.5 × 10^−2^ μM following P_2_X_4_ stimulation.

### 3.2 Cytosolic Ca^2+^-dependent signaling

Critical to relating PM Ca^2+^ influxes to intracellular Ca^2+^ was the incorporation of a Ca^2+^ buffering term in the model. P_2_X activation can contribute millimolar total Ca^2+^ to the cytosol (see Fig. 3A) total Ca^2+^ to the cytosol depending on the duration of ATPexposure, though free Ca^2+^ was found to generally increase by less than 5 × 10^-2^pM in Hide *et al.*[8], against which we fit our model. We therefore utilized the ratiometric measurements of Fura-loaded cells presented in [8] to estimate the free Ca^2+^ following substantial buffering by Ca^2+^ binding proteins [23]. Accordingly, we plot our predictions of Ca^2+^ transients following P_2_X stimulation in Fig. 3. To obtain reasonable matches with the Hide et *al.*measurements, refitting of the P_2_X constants was required. Overall, we were able to reasonably reproduce Ca^2+^ transients measured following 100 μM and 1 mM ATP treatments, though our predicted currents at 100 μM ATP decayed more rapidly than indicated by the Ca^2+^ transient data. At 10 μM ATP, however, we predicted much smaller Ca^2+^ transients, as our model suggests only P_2_X_4_ significantly contributes at this concentration. It is possible that there may be cell-to-cell variations in the expression of P_2_X_4_ and P_2_X_7_ that could account for this unexpected behavior, or the activation of a purinergic receptor not accounted for in our model.

Given our fitted model, integration of the P_2_X current profile over 400 seconds and noting roughly 8% of P_2_X current is attributed to Ca^2+^ [6], we estimate that the channels conduct sufficient Ca^2+^ to elevate the cytosol concentration by several millimolar. However, since analogous ATP exposure profiles in Fig. 3 reflected roughly 50 nM increases in Ca^2+^ content from basal levels, the vast majority of Ca^2+^ that enters the cytosol is buffered, as is typical in a variety of eukaryotic cells [23]. Therefore, it is evident that Ca^2+^ is significantly buffered in microglia, although we are unable to attribute the buffering to a specific set of Ca^2+^ proteins. Two candidates for buffering include CaM and ionized calcium-binding adapter molecule 1 (Iba-1), in addition to contributions from binding to the plasma membrane [44] and Ca^2+^ indicators used for characterizing the cytosolic Ca^2+^ transients. Although CaM content may be scarce in resting microglia[45], the ability to activate NFAT suggests there exists an extant pool capable of binding Ca^2+^. Further, Iba-1 is a known Ca^2+^-binding protein in microglia that could contribute to the buffering of cytosolic Ca^2+^ [46].

In addition to Ca^2+^ influxes and effluxes mediated by NCX and P_2_X, we reflect Ca^2+^ exchange with the endoplasmic reticulum (ER). Specifically, we include endoplasmic reticulum (ER) Ca^2+^ uptake via SERCA as well as a concentration-dependent ER Ca^2+^ leak in order to maintain constant ER Ca^2+^ content of 800 μM[12]. To our knowledge measurements of SERCA function and effects on ER Ca^2+^ in microglia are not well established, therefore we utilized SERCA parameters from Shannon *et al.*[21]. In principle, the ER Ca^2+^ leak assumed in our model to counterbalance SERCA uptake likely corresponds to spontaneous IP_3_-mediated ER release [12], though including such contributions was beyond the scope of our study, for which we assumed P_2_Y are inhibited. In Fig. S5 we demonstrate that ER Ca^2+^ load increases in response to P_2_X activation with a modest lag relative to the cytosolic Ca^2+^ transient, but returns to resting load after the termination of the membrane current. In other words, ER Ca^2+^ load is increased only transiently, which indirectly suggests that Ca^2+^ entering through the PM is extruded by another mechanism, namely NCX exchange. Hence, physiologically normal rates of Ca^2+^ entry via plasma membrane Ca^2+^ influx were compensated by NCX extrusion, and therefore were necessary for maintaining ER Ca^2+^ at resting levels when subject to infrequent or low-amplitude P_2_X_4_ stimulation.

**Figure 3:**
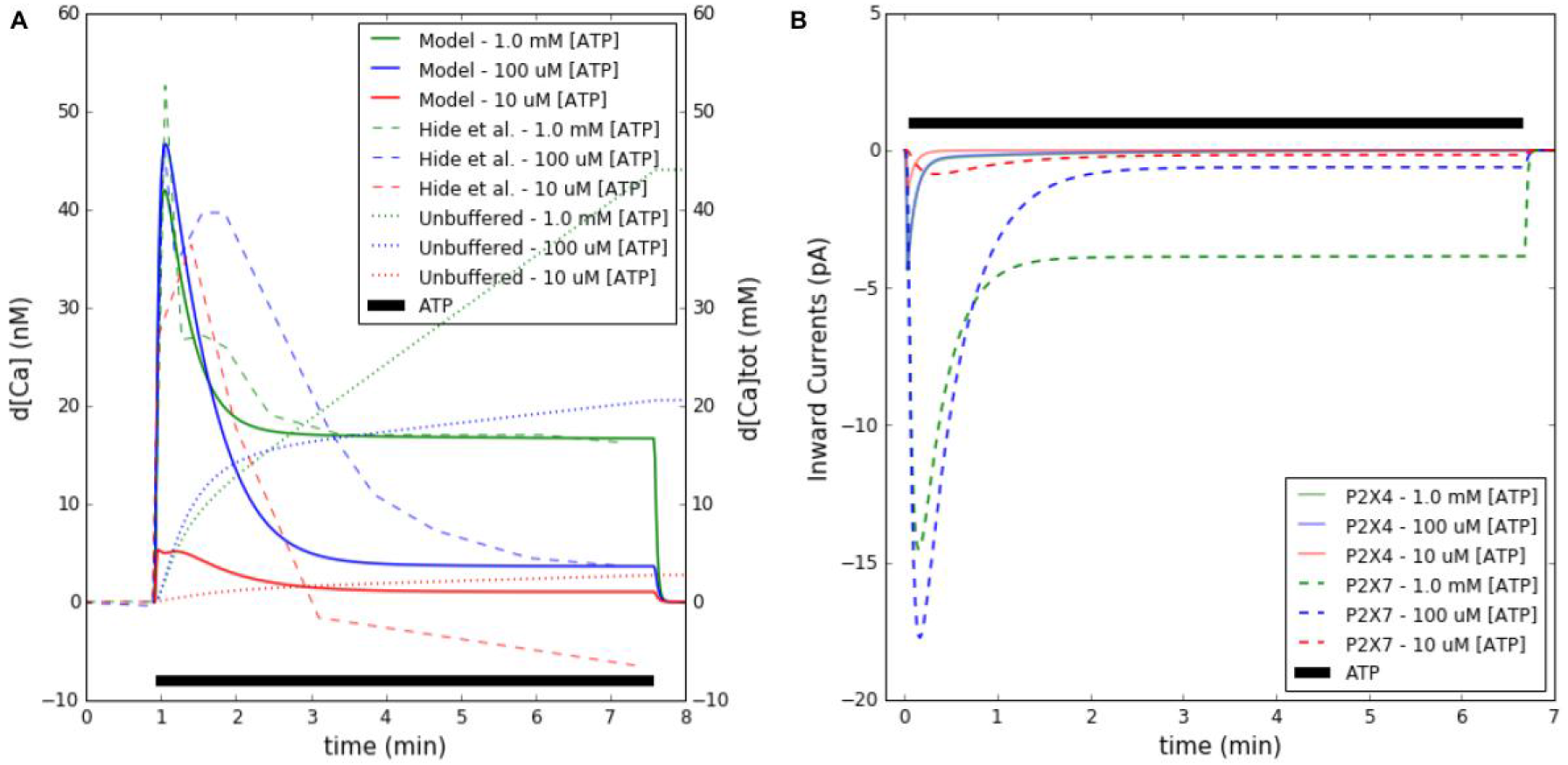
A. Predicted free cytosolic Ca^2+^ (solid) with respect to time, subject to ATP treatments of 1.0 × 10^1^ μM-1.0mMATPfor 400 s (black), compared to experimental data from mouse microglia (dashed) [8] Complementary ER Ca^2+^ transients are reported in Fig. S5. B. Contributions of P_2_X_4_ and P_2_X_7_ currents to the Ca^2+^ transients. Parameters and conditions to model the experimental conditions are listed in Table S2.

Ca^2+^dependent NFAT activation constitutes a prominent signal transduction pathway implicated in the release of inflammatory agents [47, 48] including TNF*α*[8, 27,49,50]. Of these, Ca^2+^-binding to CaM and CN, as well as subsequent binding of the Ca^2+^-saturated proteins to NFAT, is an established substrate for activation of this pathway [15]. Our implementation of the CaM/CN model was originally developed for cardiomyocytes by Saucerman et al [20], though we utilized parameters adapted by Bazzazi *et al.* [22] for HEK cells. Given that time-dependent CN activation data are not available in glia, we sought qualitative agreement between measurements of CN activation by Bazzazi *et al.*, subject to Ca^2+^ transients that we tuned to be similar in amplitude to their data. Fig. S4 summarizes our qualitative comparisons, namely that the predicted CaM/Ca^2+^-saturated CN at low stimulation frequencies reaches a maximum that lags the peak Ca^2+^ concentration by several seconds, while its decay is less rapid than that of the Ca^2+^ transient. Further, a rapid frequency of pulsatile ATP exposure (1 second pulses at 0.5 Hz) was sufficient to maintain an elevated CaM/Ca^2+^-saturated CN. Both trends were consistent with the HEK data from Bazzazi *et al.*, though our data reflect slower relaxation kinetics, which we attribute to different rates of Ca^2+^ uptake and extrusion mechanisms in microglia. Hence, we anticipate that transient increases in Ca^2+^ arising from brief P_2_X stimulation will not lead to sustained CN activation, in contrast to transients of sustained duration.

We next tuned the kinetics of NFAT activation in our microglial model to agree with Western blot-based measurements of activated (dephosphorylated) NFAT from [51], for which the dephosphorylated factor was reported for several 3 mM ATP exposure durations and concentrations (15 min). Notably, we adjusted most of the reaction constants associated with NFAT cycle model from Cooling *et al.*[15] to reproduce percent changes in NFAT with respect to ATP exposure time up to 60 minutes, in accordance with the microglial data from Ferrari *et al.*[17].

Our predictions shown in Fig. 4 are in good agreement with experimental data as exposure time was varied, and for variations of concentrations up until 3 mM ATP. A decline in activated NFAT was reported by Ferrari *et al.* that we could not capture in the model. Moreover, assuming that NFAT activation at ATP concentrations below 1 mM arise from P_2_X_4_-mediated Ca^2+^ current and P_2_X_7_ increasing contributes Ca^2+^ at and above this concentration, both channels are capable of eliciting NFAT responses.

**Figure 4:**
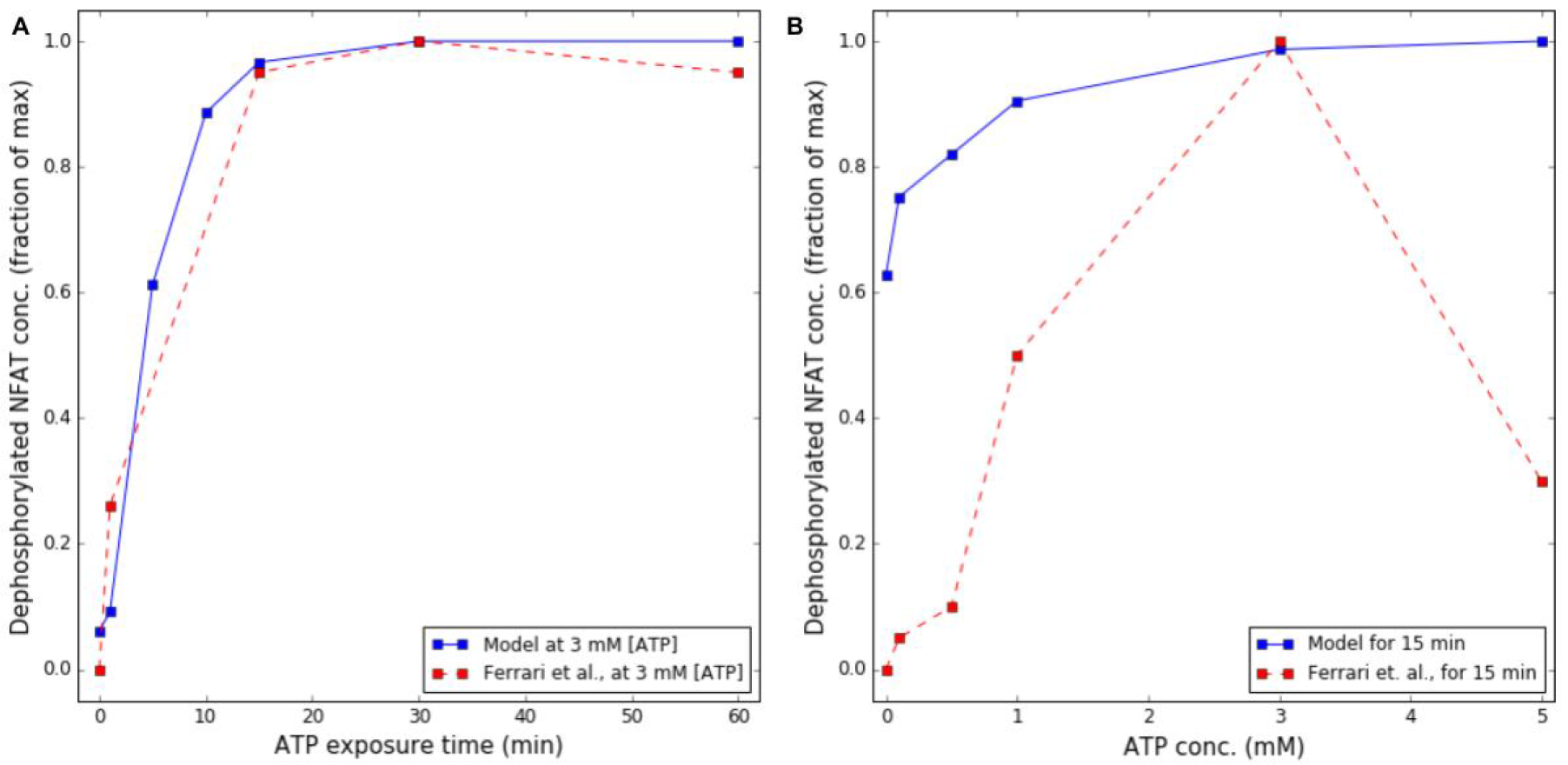
A. Activated (dephosphorylated) NFAT following 3 mM ATP treatment for up to 60 minutes recorded in Ferrari *et al.* [51](dashed) and predicted by the model (solid). B. Analogous predictions of activated NFAT, subject to 15 minute treatments of ATP up to 5 mM versus experiment [51] Parameters and conditions to model the experimental conditions are listed in Table S2.

### 3.3 Transcriptional regulation

P_2_X activation, followed by elevation of cytosolic Ca^2+^ load whereby NFAT and p38 are activated (see Sect. A.5.1), culminate in cytokine production [8, 3, 52]. We therefore utilized phenomenological models to couple NFAT and pp38 control of TNF*α* transcription, as summarized in Sect. A.3.5. Specifically, we assumed that TNF*α* mRNA transcription is driven by the formation of a NFAT/DNA complex[53], the transcribed mRNA is partially protected from degradation in a pp38-dependent manner [54], thereby permitting increased rates of TNF*α* protein translation and exocytosis [55]. It is important to note that additional regulatory mechanisms ultimately control the exocytosis of molecular cargo [56], but are represented only by approximation in this study.

In Hide *et al.* [8], enzyme-linked immunosorbent assay (ELISA) assays were conducted both 3 hours after 0 mM to 1 mM ATP treatments, as well as at 1 hour intervals using 3 mM ATP. The predicted TNF*α* production rate data were fitted to experimental data provided by Hide *et al.* (see Fig. 5), primarily through modulating parameters that control messenger ribonucleic acid (mRNA) production, its degradation, translation and P_2_X_7_-dependent exocytosis. Here we make a simplifying assumption that the TNF*α* exocytosis measured via Hide *et al.* scales proportionally to the mRNA production rate, provided that P_2_X_7_ is activated, as we do not yet have a precise model for regulatory control of the entire exocytosis process following NFAT activation. The predicted TNF*α* production data positively correlate with ATP concentration and duration of exposure, although the model overestimates release rates at 3 mM ATP and is less sensitive to ATP concentration than reported by Hide *et al.*. Though further refinement of the TNF*α* transcription model could improve agreement with experiment (see Sect. 4.6), our model nevertheless demonstrates that even low levels of ATP can trigger TNF*α* production, thereby suggesting sensitivity to both P_2_X_7_ and P_2_X_4_ activation.

**Figure 5:**
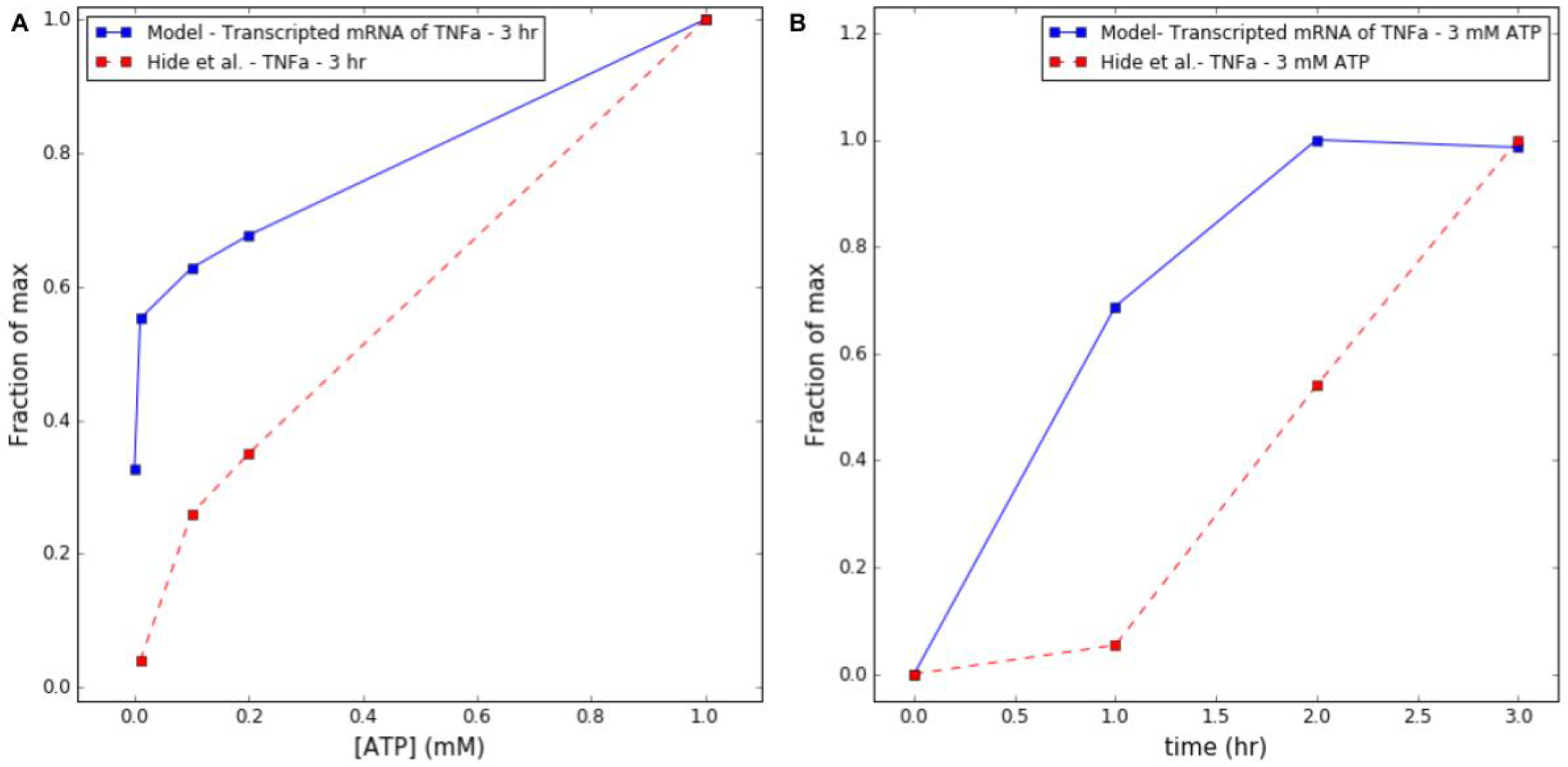
A. Predictions (solid) of TNF*α* mRNA production following 3 hours of micromolar to 1 millimolar ATP exposure versus exocytosed protein collected by Hide *et al.*[8] (dashed), assuming that exocytosed TNF*α* scales proportionally to mRNA transcription. B. TNF*α* release predictions (solid) as a function of time (hours), following 3 mM ATP treatment versus experiment (dashed) Parameters and conditions to model the experimental conditions are listed in Table S2.

### 3.4 Effects of ATP waveform on intracellular Ca^2+^ dynamics and TNF*α* production

Our overarching hypothesis in this study was that the duration, frequency and amplitude of ATP stimulation control the dynamics of the intracellular Ca^2+^ transient and ultimately, the ability to trigger TNF*α* production. This hypothesis was chiefly motivated by observations that 1) low micromolar, high frequency (up to 0.5 Hz) quanta of ATP are released by astrocytes as part of their physiological function [57], 2) P_2_X_4_ and P_2_X_7_ present markedly different ATP-sensitivities and deactivation kinetics [14], and 3) non-P_2_X-specific Ca^2+^ entry via ionophores induces TNF*α* mRNA production in T cells [58] and CCL3 (a chemokine) mRNA via NFAT in microglia [47]. To test this hypothesis, we subjected our model validated in the previous section to ATP pulses ranging in duration from 1-19 seconds, amplitudes of 0-1 mM, and frequencies of 0.083-0.5Hz. Here, we first discuss the degree to which these conditions changed the average intracellular Ca^2+^ content from values maintained at rest, after which we phenomenologically relate the Ca^2+^ content to the propensity for TNF*α* release. Based on data presented in Fig. 6A, we find that high frequencies of low amplitude ATP stimulation are sufficient to increase the cytosolic Ca^2+^ load in a manner sufficient to promote TNF*α* mRNA production above resting levels, although to a lesser extent than changes following millimolar ATP concentrations. Given that P_2_X_4_ responds to micromolar ATP, it is apparent that P_2_X_4_ contributes most significantly under high frequency ATP stimulation, whereas P_2_X_7_ accounts for the majority of Ca^2+^ entry at high ATP concentrations. We note that a report [49] demonstrated that TNF*α* release was not reported in P_2_X_7_ knock-out microglia under 24 hours of ATP treatment [49]. An important distinction in our protocol, however, is that we utilize short bursts of ATP that reduce that rate of P_2_X_4_ desensitization.

Second, we demonstrate in Fig. 6B rates of ATP-mediated TNF*α* production under conditions analogous to those in panel A. Importantly, we observe 1) significant rates of TNF*α* production under high frequency, low amplitude ATP stimulation and 2) that the TNF*α* production rates do not strictly correlate with increase Ca^2+^ load, as TNF*α* production rates decrease with decreasing frequency. Overall, these results appear to establish bounds for the frequency and amplitude capable of evoking transcriptional responses.

**Figure 6:**
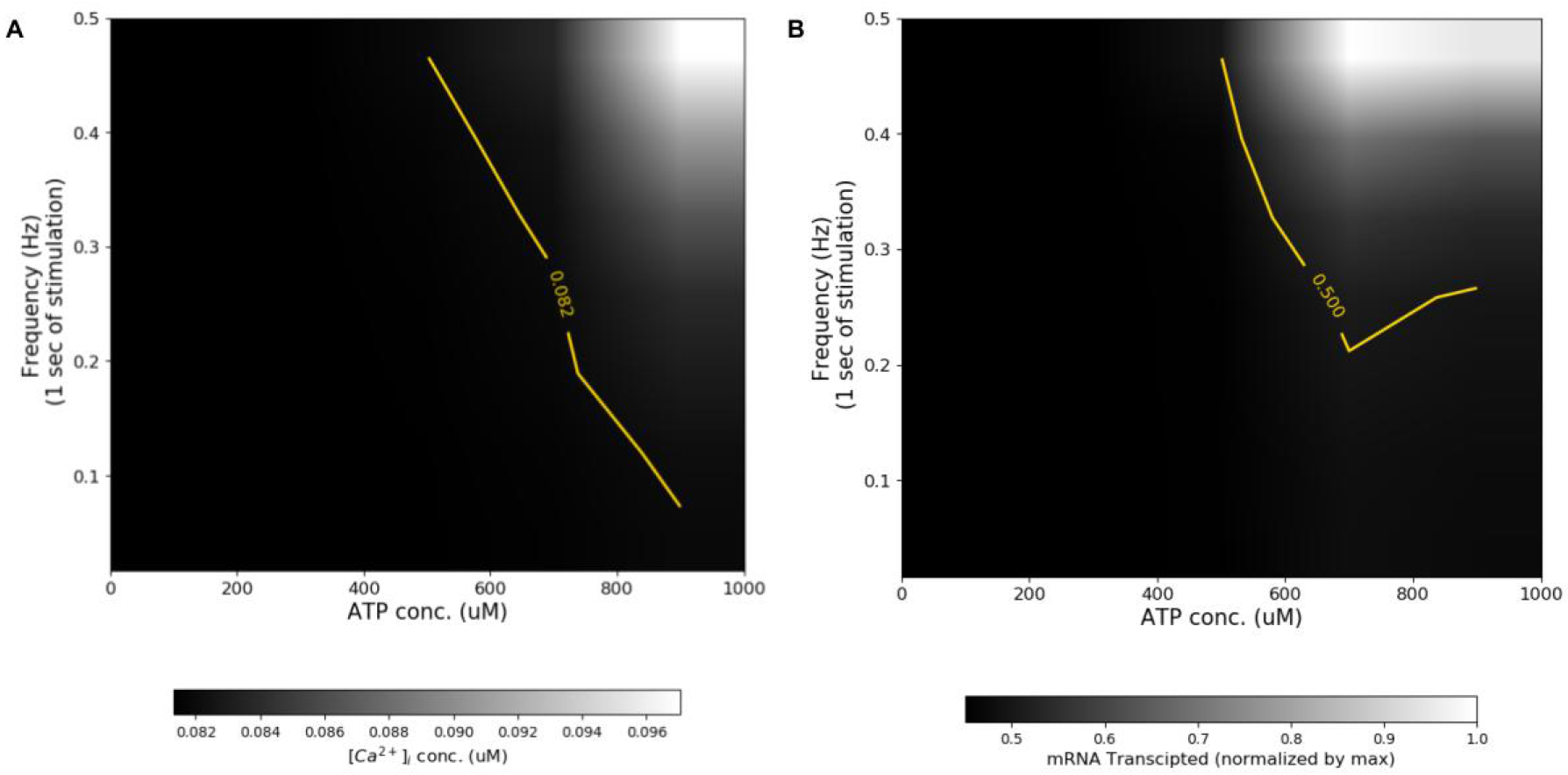
Predictions of free cytosolic Ca^2+^ (A) and TNF*α* production (B) following pulsatile (1 second duration) ATP exposure for 100 seconds. ATP pulse frequency is varied from 0-0.5 Hz and concentration is varied up to 1 mM. The TNF*α* data were normalized by the maximum TNF*α* production under conditions considered. For reference, thresholds are drawn at Ca^2+^=8.1 × 10^−2^ μM and TNF*α*=0.6. Ca^2+^ transients are illustrated in Fig. S6.

### 3.5 Sensitivity analysis and effects of partial (ant)agonism on TNF*α* production

Lastly, given that TNF*α* transcription and release is collectively controlled by several Ca^2+^-dependent compounds in our microglial model, we utilize sensitivity analysis to highlight parameters that most significantly influence the model predictions. Our central goal here was to provide predictions that can be tested with common experimental protocols, such as controlling gene expression, or the application of protein-specific inhibitors and agonists. Here, protein concentration was varied as a surrogate for up-or down-regulation, or to a certain extent, partial inhibition or agonism of a given target. In Fig. 7, we report the sensitivity of TNF*α* mRNA production as a function of plasma membrane Ca^2+^ entry, as determined by purinoceptor concentration, NCX Ca^2+^ flux, and the rate of inward Ca^2+^leak, subject to low and high ATP concentrations. Here, each line represents the rate of TNF*α* mRNA production, *r_mRNA_* relative to control versus the channel expression relative to control [*P*]. The slope of a given line, Δr_mRNA_/Δ[*P*], indicates the sensitivity of the mRNA production relative to changes in channel expression. Since the P_2_X receptors and the Ca^2+^ leak contribute inward Ca^2+^ current, TNF*α* production is positively correlated with the concentration (P_2_X) or rate (leak) relative to control values. Importantly, the slope increases with increasing channel expression, which suggests that greater rates of TNF*α* production are realized relative to control or equivalently, mRNA production could occur at lower ATP concentrations. Similarly, higher rates of Ca^2+^ influx via a non-specific, inward Ca^2+^ leak sensitizes TNF*α* mRNA production, hence, the threshold for TNF*α* production via P_2_X stimulation is reduced. Analogously, reducing extracellular Na^+^ decreases and potentially reverses the rate of Ca^2+^ extrusion, which has the net effect of increasing intracellular Ca^2+^ relative to control. Hence, decreased rates of Ca^2+^ extrusion that arise from decreasing Na^+^ sensitizes TNF*α* mRNA production.

In Fig. 8 we report similar measures of TNF*α* production sensitivity to variations in CaM, CN, and NFAT. Given that the Ca^2+^-dependent activation of these model components promote TNF*α* expression, decreases in concentrations of these protein depressed TNF*α* production, and conversely, their increase accelerated TNF*α* production rates. Of these, TNF*α* production was most sensitive to the modulation of CaM and CN, with a relatively minor dependence on ATP concentration. Among the proteins considered, CaM is known to increase expression upon activation [45], which suggests that TNF*α* production may be accelerated in activated microglia.

**Figure 7:**
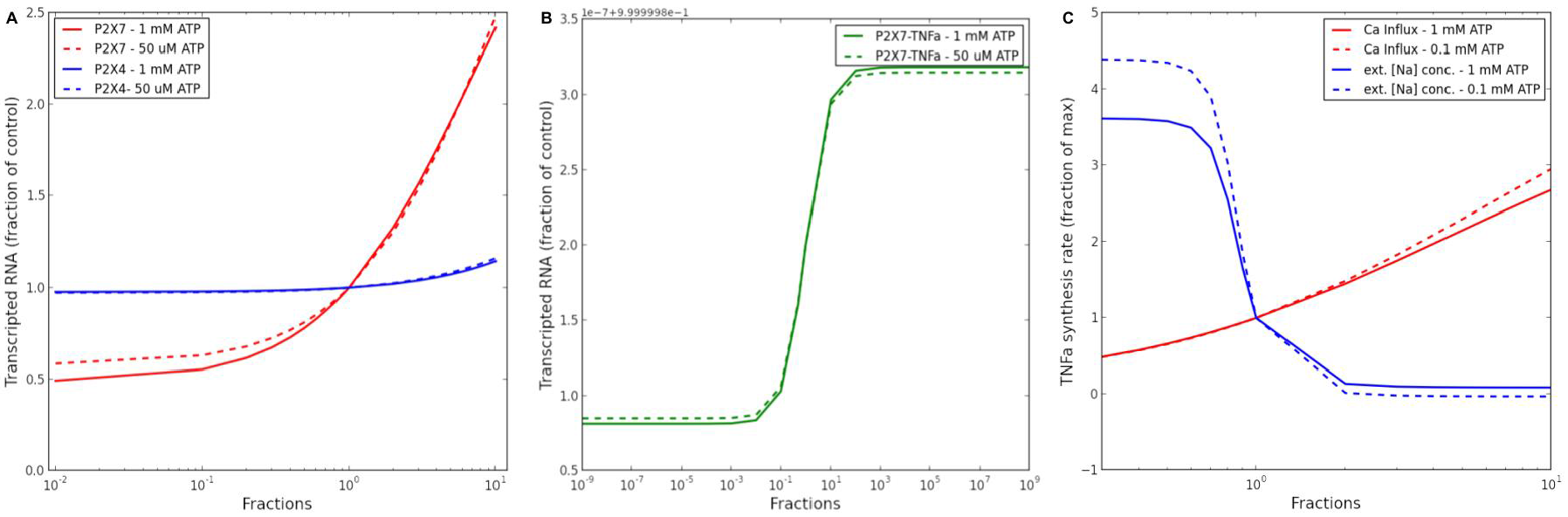
A. Sensitivity of TNF*α* production as a function of P_2_X_4_ (red) and P_2_X_7_ (blue) expression, subject to conditions from Fig. 8. B. Sensitivity analysis of TNF*α* dependency on P_2_X_7_ open state. C. Analogous predictions for changes in inward Ca^2+^ leak rate (blue) and extracellular Na^+^. Reduction of Na+ reduces the Ca^2+^-extrusion rate from forward-mode NCX activity.

**Figure 8:**
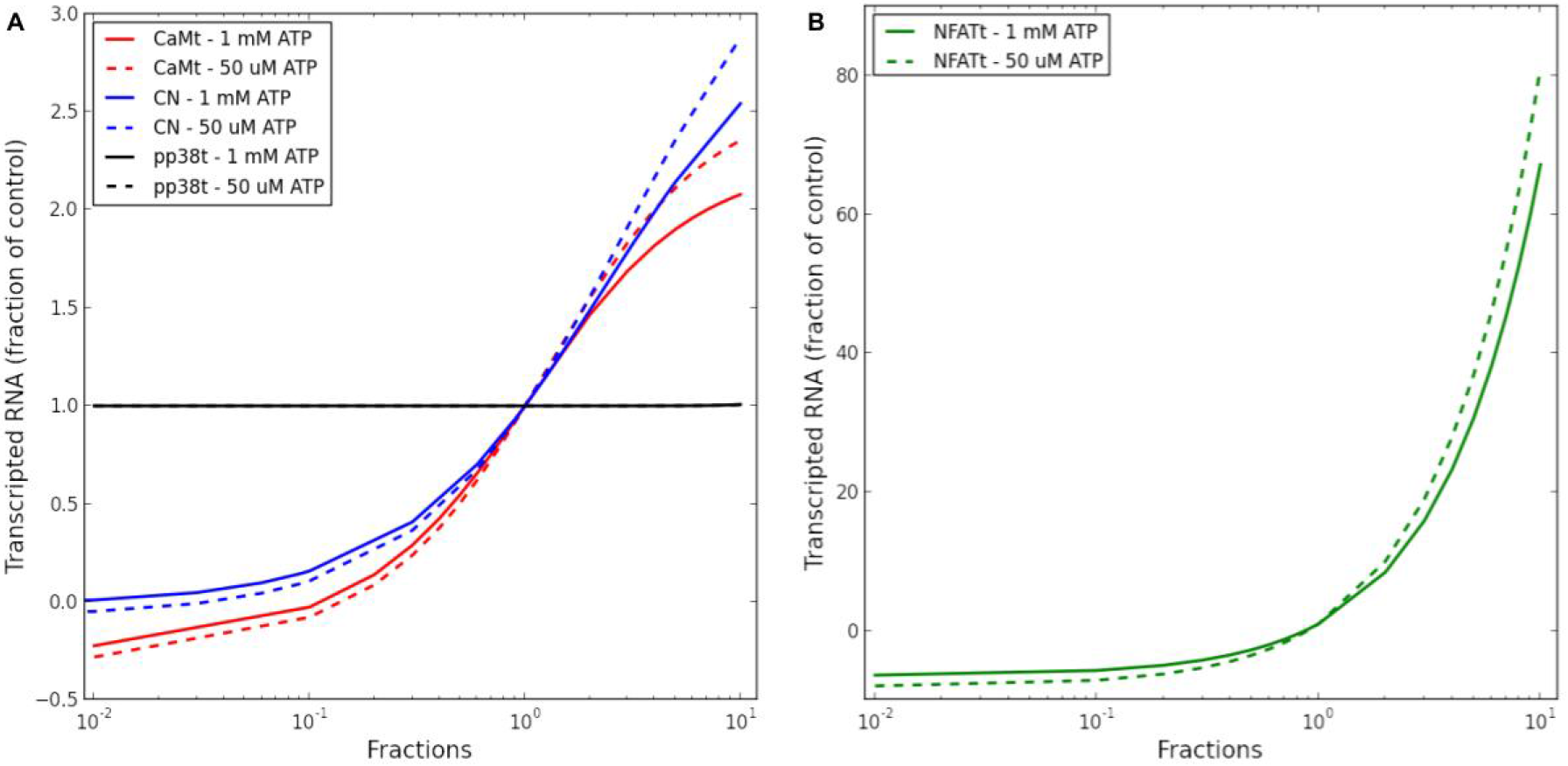
Sensitivity of predicted synthesis rate of TNF*α* to variations in model parameters. Degree of TNF*α* synthesized as a function of total CaM(red), CN(blue), NFAT (green) (B) or pp38 (black) concentration with respect to 5 × 10^−1^ Hz 0.05 and 1.0 mM ATPpulses for 100 seconds.

## 4 Discussion

### 4.1 Overarching outcomes

In this study, we have established a predictive, computational model The computational model recapitulates ATP-dependent activity profiles for two prominent P_2_X channels, parallel rises in intracellular Ca^2+^, activation of the Ca^2+^-dependent NFAT pathway, and ultimately cytokine production. With this minimal model, we assess the extent to which the intracellular signaling cascades are coupled to characteristics of extracellular ATP exposure, including the amplitude, frequency and duration of stimulation, as well as the sensitivity of activating signal transduction to variations in protein expression. The breadth of ATP signatures considered were intended to be representative of transient, small amplitude release events within synaptic junctions commonly surveilled by microglia [59, 27], as well as prolonged, concentrated ATP pools that could emerge from neuron damage [60]. The variations considered in the sensitivity analyses were reflective of known phenotypical changes in expression during microglial activation, as well as pharmacological or genetic strategies used in a variety of experiments to manipulate and understand microglia physiology. The model we developed integrates both biophysically motivated and phenomenological descriptions of purinergic receptors, NFAT-mediated gene expression pathways, cytosolic Ca^2+^ handling, as well as provides a framework to link a wide body of experiments probing microglial function (reviewed here [3, 61]). With this model we characterized the relative contributions of P_2_X_4_ and P_2_X_7_ in driving TNF*α* production across a broad range of ATP time and concentration profiles and demonstrated how changes in protein expression and activity typical of microglial activation tune TNF*α* mRNA expression. Most significantly, we provide computational evidence that TNF*α* mRNA expression is possible from transient, high frequency exposure to ATP signals, thresholds for mRNA production are reduced under conditions typical of activated microglia, e.g. increased P_2_X and CaM expression.

### 4.2 P_2_X receptor activation

Recently, there has been considerable focus on the modulation of purinergic receptors, including P_2_X_4_ and P_2_X_7_, in controlling pain and inflammation, as reviewed in [62]. Both P_2_X receptors are responsive to ATP released from damaged cells[59, 63, 64]. Although both channels are activated by ATP, the micromolar versus millimolar affinities of P_2_X_4_ and P_2_X_7_, respectively, raise the possibility that the channels have distinctive roles in their responses to extracellular agonists.

Toward addressing this hypothesis, we implemented state-based models of P_2_X_7_ and P_2_X_4_ activation that could predict channel open probabilities over a broad ranges of ATP exposures. In turn, the current profiles were used to predict rates of intracellular Ca^2+^ entry. The channel models considered in this study were developed in prior works that examined P_2_X channel activity for endogenous ATP, as well as several pharmacologic agonists and antagonists [37, 5,19, 39]. These models consisted of at least eight and as many as 32 states to capture nuances of desensitization and changes in channel gating arising from pharmacological treatment. Our primary objective was to exclusively examine channel responses to ATP over a range of concentrations, durations, and frequencies, thus we opted for a simplified model with fewer free parameters. Overall, the reduced models, which shared the same number of states but with different gating parameters, reproduced experimentally-measured and predicted current profiles with acceptable accuracy. For this reason, the parsimonious model was appropriate for predicting increases in intracellular Ca^2+^ load, assuming that 8% of each channel’s predicted current corresponded to Ca^2+^ influx, as suggested by Garcia-Guzman *et al.*[6]. We note, however, that further refinement of the model to reflect responses to P_2_X_7_ and P_2_X_4_ agonists like 3′-O-(4-Benzoyl)benzoyl ATP (BzATP) and ivermectin, respectively, could necessitate additional gating terms described in Yan et *al.* and Zemkova *et al.* [37,19]

To determine the extent to which P_2_X_4_ modulation impacted steady-state Ca^2+^ load and subsequent TNF*α* production, we utilized a basic sensitivity analyses approach described in [28]. Our predictions of P_2_X_4_ activity indicate that low amplitude ATP exposures of short duration (< 30 seconds) across a broad range of frequencies were sufficient to trigger significant Ca^2+^ entry. Microglial P_2_X_4_ thus, could serve a predominant role in processing small, highly transient, ATP signals arising from normal communication between in neural synapses [59], to activate a microglia-dependent modulation of synaptic function. An exciting possibility that remains to be tested is whether there are lower thresholds for ATP concentration and exposure duration that is sufficient to activate P_2_X_4_-dependent Ca^2+^ signaling in microglia. For instance, how many synapses with elevated ATP would a single microglia cell need to sample during a short duration (milliseconds-to-seconds) to activate a P_2_X_4_dependent Ca^2+^ response. Furthermore, since prolonged exposure to ATP is associated with increased membrane trafficking and mobility of P_2_X_4_ [9], higher P_2_X_4_ activity could prime microglia to have increased sensitivity to ATP, thus lowering the threshold of ATP needed to induce P_2_X_4_-dependent Ca^2+^ responses. The sensitivity analyses we conducted indeed demonstrate greater propensities for TNF*α* mRNA production under conditions of ten-fold higher P_2_X_4_ akin to activated microglia. It remains to be determined experimentally the extent to which simultaneous changes in microglial Ca^2+^ handling upon activation, such as increased CaM [45] and Iba-1 expression [65], could compensate for larger inward Ca^2+^ currents via increased P_2_X_4_ expression. Beyond changes in expression, differences in subcellular localization could additionally counterbalance the effects of increased P_2_X_4_ activity, such as the asymmetric distribution and mobility of P_2_X_4_ between the microglial soma, and the microglial processes [9], as well as the localization of ATP encountered presumably by the microglia process at the synapse. Careful characterization of P_2_X_4_ sub-cellular distribution, thus, may be important for understanding the channels ability to prime microglia to increased sensitivity to ATP.

Our computational simulations indicate P_2_X_7_ responses predominate over P_2_X_4_ at high ATP concentrations and prolonged exposure durations. Our computational sensitivity analyses are also in agreement with the role of P_2_X_7_ in ATP-dependent TNF*α* production from microglia [8], and the effects of selective inhibitors of P_2_X_7_ like KHG26792 [66, 67] that suppress TNF*α* production. The distinct channel activation profiles of P_2_X_4_ and P_2_X_7_ likely provide microglia the ability to responded differently to low or high levels of ATP, by activating a neurotrophic [68], or cytokine response respectively. Moreover, selective inhibitors of P_2_X_7_ may be useful in the case of pathologically elevated ATP, such as that seen after a traumatic brain injury, while selective agonists or antagonist of P_2_X_4_ may be more useful in modulating microglia when subject to physiological levels of ATP.

### 4.3 Maintenance of cytosolic Ca^2+^ content

Activation of microglia and exocytosis of cytokines and chemokines are strongly correlated with increased intracellular Ca^2+^ content. We observed that the sensitivity of ATP-triggered TNF*α* production increased at elevated Ca^2+^ inward leak rates (see Fig. 7). The increased cytosolic Ca^2+^ load can arise from an arbitrary source, such as Ca^2+^ selective ionophores, which can increase mRNA expression independent of toll-like receptor signaling or purinergic receptor activation [47]. A primarily goal of our model was therefore to couple the Ca^2+^ current carried by activated P_2_X receptors to the intracellular Ca^2+^ pool. Our model found that cytosolic Ca^2+^ is strongly buffered, with as much as 90% of the inward Ca^2+^ buffered within the cytosol. CaM, CN and Iba-1 are among the most prominent Ca^2+^-binding proteins identified in microglia, [3, 7, 13]. However, it is no clear if the microglia express sufficient CaM, CN and Iba-1 to account for 90% buffering of cytosolic Ca^2+^There may be other cytosolic Ca^2+^ proteins or molecules that we do not account for, or non-specific binding of Ca^2+^ to negatively-charged membrane that may explain this discrepancy. In cardiac tissue, of the intracellular components in common with microglia, ATP and SERCA buffer a substantial fraction of Ca^2+^ at rest and following Ca^2+^ channel activation, with SERCA accounting for the majority of Ca^2+^ binding [69]. Similarly, it is apparent that low-(micromolar) and high-affinity (submicromolar) Ca^2+^ binding to phospholipid membranes contribute roughly 42 and 15 μM of buffering capacity [44]. Given the large surface area to volume ratio in resting microglia [70], it is plausible that its plasma membrane could significantly contribute to buffering intracellular Ca^2+^.

We additionally observed that the sensitivity of ATP-triggered TNF*α* production increased at elevated Ca^2+^ inward leak rates (see Fig. 7). This behavior is analogous to the positive correlation between TNF*α* production sensitivity and P_2_X expression, except here the change in cytosolic Ca^2+^ load can arise from an arbitrary source, such as treatment of microglial cells with Ca^2+^ selective [47] or non-selective ionophores [71]. In this regard and to a certain extent, diverse mechanisms that increase cytosolic Ca^2+^ load may comprise a positive feedback process, as the Ca^2+^-dependent activation of genes that initially promote autocrinic activity or enhance Ca^2+^ channel current in principle can promote further gene expression. This feedback could provide a basis for an integrative activation of immune responses [49], whereby both danger-associated and pathogen-associated molecular patterns must be presented to invoke an immune activity. However, such feedback is nuanced, as it has been demonstrated that pathogen mimetic lipopolysaccharides (LPS) treatment blunts ATP-mediated P_2_X responses [71], which warrants further model development to investigate this hypothesis.

In general, PM Ca^2+^ currents are counterbalanced by Ca^2+^ extrusion mechanisms to maintain constant Ca^2+^ load at rest. In our study, modest levels of Ca^2+^ entry via P_2_X activation were compensated by forward mode extrusion of intracellular Ca^2+^ via NCX. Given the slower rate of Ca^2+^ efflux via NCX relative to SERCA uptake, transient increases in ER Ca^2+^ load were apparent in our model (see Fig. S5). While IP_3_ receptor-mediated Ca^2+^ release was beyond the scope of this study, it is possible that the transient increases in ER Ca^2+^ load could lead to more frequent spontaneous or IP_3_-triggered ER Ca^2+^ release events [3, 7]. IP_3_-triggered ER Ca^2+^ release could lead to localized regions wherein Ca^2+^ is strongly elevated relative to bulk cytosol, which could disproportionally influence Ca^2+^ signal transduction pathways. Future models that account for the microglial morphology and spatial distribution of important signaling proteins may provide additional insights in microglial function.

### 4.4 Ca^2+^/CaM/CN/NFAT signal transduction

In microglia, CaM is linked to cytoskeletal remodeling [72] and the activation of CN[16]. The CN phosphatase is well-established to play a significant role in glial physiological function, as reviewed in [16]. In response to elevated cytosolic Ca^2+^ levels, CaM and CN activation ensues whereby NFAT is dephosphorylated. The dephosphorylated NFAT translocates to the nucleus, where it modulates transcription. Similar levels of NFAT activation are reported in microglia treated with Ca^2+^ ionophore (A23187) [47], which suggests that Ca^2+^ alone is sufficient to invoke this response. CN and NFAT activation are known to be dependent on stimulation frequency [22, 15], thus it is of interest to understand the relationship of NFAT driven TNF*α* expression in response to transient and sustained ATP stimuli. In this study, we specifically investigated the coupling between CN activation and dephosphorylation of NFAT, for which the latter is a substrate for TNF*α* transcription [8]. We assumed in our model that full activation of CN proceeds through binding of Ca^2+^saturated CaM, as described in Saucerman et al.[20]. We found that rates of increased TNF*α* directly correlated with the CaM-bound CN concentration and NFAT activation. Moreover, high frequency P_2_X stimulation that led to sustained increases in cytosolic Ca^2+^ load were sufficient to maintain CN in an activated state capable of driving NFAT-dependent TNF*α* production. Low frequency P_2_X_7_ stimulation exhibited analogous behavior, as our computational model showed that inhibition of NFAT by nearly 100% substantially suppressed the TNF*α* mRNA production rate 1 mM ATP, consistent with reported reductions in TNF*α* following treatment with the NFAT inhibitor VIVIT in [18, 73]. Furthermore, our computational model showed similar reductions when inhibiting CN activity, which were comparable to reductions reported in Rojanathammanee *et al.* [73] following 1 μM treatments of the CN inhibitors FK506. On one hand, the ability of the computational model to capture both the Ca^2+^-dependent activation of NFAT and the blunting of TNF*α* production by CaM, CN or NFAT inhibition exemplify the exquisite coupling of gene expression in microglia to intracellular Ca^2+^ homeostasis, in addition to establishing the accuracy of the model. However, it is clear that additional regulatory mechanisms convolute this coupling, as pharmacologic inhibition of CN and VIVIT does not completely suppress TNF*α* expression [18, 73].

### 4.5 ATP-triggered TNF*α* synthesis and release

Our study focused on a representative cytokine associated with an activated microglial state. Our model for TNF*α* expression was phenomenological in form, but captured key stages of mRNA transcription, with phenomenological representations for translation and Ca^2+^-dependent exocytosis. We assumed deterministic rate expressions for these stages, which were found to qualitatively reproduce TNF*α* ELISA assays following microglia activation [8, 73, 74] including following treatments with pp38, CN, and NFAT inhibitors, as well as increased mRNA expression upon increasing PM Ca^2+^ leak in analogy to ionophore treatment [47]. The agreement between the deterministic models assumed here and the experimental measurements suggests that the modeled Ca^2+^-dependent processes are abundant in number and are frequent. In counter example, we anticipate that measurements performed on an individual cell or perhaps sub-cellular basis would be fewer and less frequent, which advocate for stochastic approaches commonly used for modeling gene expression in other cell types [75].

Our model assumed that TNF*α* mRNA transcription is proportional to the nuclear content of dephosphorylated NFAT. While it is clear from our analyses that the frequency and concentration of ATP application could modulate NFAT activation and TNF*α* mRNA transcription, the apparent dependence of TNF*α* translation and exocytosis on P_2_X_7_-specific activation suggests that parallel regulatory mechanisms are required for microglial immune responses. More generally, it has been suggested that microglial activation is not ‘all or none’ and depends profoundly on the context of the activating stimuli [27]. Interestingly, TNF*α* is known to feedback on TNF*α*-receptors localized to the microglia plasma membrane, thereby contributing to further TNF*α* biosynthesis and cytokine release [76]. It is plausible that transient ATP stimuli lead to brief P_2_X activation intervals that result in a limited exocytosed TNF*α* pool insufficient for autocrinic activity, whereas prolonged ATP exposure could promote sufficient TNF*α* release to active TNF*α*-receptors. The former scenario may be appropriate for regulating synaptic plasticity mediated by TNF*α*[59], especially given the quantile nature of ATP release exhibited by astrocytes [57]. In the latter scenario, however, released cytokines could lead to more robust activation of microglial cytokine receptors, thereby driving phenotypical changes that may represent the tipping point between acute and sustained microglial activation [76].

### 4.6 Limitations

Our computational model used a minimal basis for linking microglial P_2_X activation with TNF*α* production. The model provides a starting point for adding in additional factors and fine tuning the model with new experimental data. Additional factors that could be considered in the model include microglial heterogeneity due to genetics, age, and disease, cultured microglia versus *in vivo* cells, as well as differences in the neuronal and glia populations in different regions of the CNS that influence the cell-to-cell interactions. Our study examined P_2_X activation given their importance in microglia activation and inflammatory responses. However, P_2_Y receptors are also highly expressed in microglia and respond to micromolar ATP concentrations [8]. Future computational models can integrate and explore the interaction of P_2_Y with P_2_X responses similarly to the way we compared P_2_X_4_ versus P_2_X_7_. Finally, beyond purinergic receptors, microglia express a myriad of other cell surface receptors, such as the toll-like receptors, involved in cytokine signaling, which we could integrate into future models.

In addition to intracellular, IP_3_ receptor-mediated Ca^2+^ production, we omitted store operated Ca2+ entry (SOCE). SOCE describes a mechanism, by which depletion of ER Ca^2+^ is sensed by Stromal interaction molecule 1 (STIM1), after which activated STIM promotes extracellular Ca^2+^ entry by binding ORAI1 [3] SOCE is prominent in microglia and is known to increase steady-state Ca^2+^ influx following prolonged ATP treatments [3]. However, under the conditions considered in our model, our predictions demonstrated minimal changes in ER Ca^2+^ load, given the absence of IP_3_ receptor-mediated release. Hence, therefore we do not anticipate contributions from SOCE that would influence the model predictions under the stimuli we considered. We note that introducing this contribution to cytosolic Ca^2+^ handling in subsequent models would be advantageous for examining how long-lived elevations in Ca^2+^ load control activated microglial responses including the release of cytokines, chemokines, nitric oxide as well as reactive oxygen species. [77]

Beyond purinergic receptors, myriad plasma membrane channels have been identified in microglia. Many of these channels, particularly those controlling potassium transmembrane currents [78], have altered expression levels depending on the maturity and potential pathological state of the microglial cell
[79]. Given the prominent role of potassium channels in determining the membrane potential, we could expect phenotype-dependent changes in the electrogenic driving forces governing voltage-gated channel and exchanger activity. For instance, though our model demonstrates NCX operates in forward mode to extrude cytosolic Ca^2+^, hyper-polarization of the membrane potential could induce reverse-mode NCX activity that would increase intracellular Ca^2+^ load. Overall, inclusion of channels that overwhelmingly control microglia membrane potentials would permit more detailed examinations of microglia activation and phenomena in pathological states.

Our model of TNF*α* expression assumed NFAT-dependent mRNA production, followed by degradation [48], translation and exocytosis process that were phenomenologically dependent on p38 phosphorylation. However, there exist a variety of regulatory mechanisms of gene expression that were not considered in this study. For instance, extracellular signal-regulated kinases (ERK) and c-Jun N-terminal kinase (JNK) inhibition in microglia has been demonstrated to both reduce CXCL2 mRNA expression and TNF*α* exocytosis [8] and following P_2_X_7_ activation [48]. Hence, it is plausible that several mitogen-activated protein kinase (MAPK)s work in tandem to finely control TNF*α* mRNA expression. Additionally, translation of expressed TNF*α* mRNA and its eventual exocytosis is tightly regulated by diverse trafficking and secretion pathways, as reviewed in Stanley *et al.* [56] for TNF*α* release in macrophages. Refinement of our model to include such pathways, as well as their differential regulation by P_2_X_7_ versus P_2_X_4_, could elucidate the cocktail of stimuli required for encoding specific patterns of microglia-derived signaling molecule expression.

## 5 Conclusions

In this study, we have developed a computational model of Ca^2+^ handling in microglia that links ATP-dependent activation of P_2_X_4_and P_2_X_7_activation to production of a representative cytokine, TNF*α*. We validated this model against a wealth of experimental data collected for these purinergic receptors and intracellular components that regulate Ca^2+^ homeostasis and gene expression. We have demonstrated using this model how the temporal signature of ATP activation influences the ability to evoke TNF*α* production. Furthermore, we examined how variation of the expression of the microglia proteins considered in this model impacts the ATP-sensitivity of TNF*α* production, so as to provide insight in how microglia respond and adapt to their environment. We anticipate that this model will be invaluable generating and testing hypotheses pertaining to microglial physiology at a systems level, as has been demonstrated in other computational models of mammalian cellular function [4].

## 6 Acknowledgements

Research reported in this publication, release was supported by the Maximizing Investigators’ Research Award (MIRA) (R35) from the National Institute of General Medical Sciences (NIGMS) of the National Institutes of Health (NIH) under grant number R35GM124977. PKH would like to acknowledge John Gensel, Ph.D. for critical discussion of the manuscript.

